# Trapping more seeing more, benchmarking a new trap column for low-input proteomics

**DOI:** 10.64898/2026.11.12.766175

**Authors:** Dennis Dannecker, Todd Blackburn, Liang Chen, Jarrod J. Sandow, Karl Mechtler, Manuel Matzinger

## Abstract

Nanoflow liquid chromatography coupled to mass spectrometry (LC-MS) has become central to low-input and single-cell proteomics (SCP), where trap-and-elute workflows offer sample loading flexibility and analytical column protection but introduce additional dead volume, surface area, and a largely uncharacterized trade-off between flush direction and chromatographic performance. Here, we present a systematic, iterative benchmarking study of a purpose-built family of NanoShield™ trap columns against the established PepMap™ Neo trap cartridge and direct injection. Using a standardized workflow built on a commercial HeLa digest standard, we evaluated performance across two mass spectrometry platforms, multiple LC gradient lengths, both flush directions, and a range of sample input amounts and injection volumes. Shortening of the NanoShield™ packed bed, from an initial 5 cm format to a 1.2 cm format, closed a substantial sample loading time penalty and largely eliminated a pronounced forward-flush performance deficit, while remaining superior to the PepMap™ in reverse-flush mode and approaching direct injection level identification numbers, peak width, and quantitative precision. A next-generation NanoShield™ featuring a stationary phase with larger pore size further narrowed the residual forward-flush penalty. These results demonstrate that trap column bed length and stationary phase chemistry are key, partially independent design levers for closing the performance gap between trap-and-elute and direct injection workflows, and provide practical, present-day guidance for trap column selection and operation in low-input proteomics.

**HIGHLIGHTS:**

- Iterative, feedback-driven benchmarking narrowed the performance gap between trap- and-elute and direct injection workflows for low-input proteomics.
- Shortening the NanoShield™ trap column from 5 cm to 1.2 cm eliminated most of the loading time penalty and forward-flush peak broadening vs the PepMap™ Neo trap cartridge.
- NanoShield™ trap column with a stationary phase featuring larger pore size (300 Å) further reduced the residual forward-flush penalty.
- In reverse-flush mode, the optimized short NanoShield™ trap column approached direct injection level IDs, peak width, and quantitative precision while retaining higher throughput.
- NanoShield™’s advantage over PepMap™ held across a range of sample inputs and injection volumes.
- Trap column length and pore size emerge as key design levers for closing the forward-flush gap, offering practical guidance for trap column selection and use.

## INTRODUCTION

Mass spectrometry (MS)-based proteomics has progressively pushed toward ever lower sample input amounts, driven by applications ranging from limited clinical biopsy material to single-cell proteomics (SCP), where total protein content can fall into the sub-nanogram^1–3^ or even low sub-picogram range^4^. At these input levels, every step of the analytical workflow, from sample preparation to chromatographic separation, carries a risk of irreversible analyte loss. Nanoflow liquid chromatography (LC) has become the standard approach for such samples because it drastically increases electrospray ionization efficiency and reduces sample dilution, thereby improving sensitivity over capillary- or micro-flow alternatives^5–7^. At the same time, nanoflow systems depend critically on minimizing post column dead volume between the separation column and the electrospray emitter, whether via a zero-dead-volume union or, more recently, an integrated pulled-tip emitter column, preserving resolving power, sharp peaks, and consequently improving sensitivity^5^.

A central architectural choice in nanoflow LC-MS workflows is whether to employ a trap-and- elute configuration, in which sample is first loaded onto a short trap column before being eluted onto the analytical separation column, or to inject the sample directly onto the analytical column. Trap-and-elute setups serve two complementary purposes: they allow sample loading and desalting at higher flow rates and larger injection volumes than would be compatible with the analytical column, and they act as a sacrificial guard column, intercepting particles and other insoluble material before these can reach and potentially damage the considerably more expensive analytical column^8–10^. This protective function, however, is not without cost. The trap column itself introduces additional surface area and dead volume into the flow path, both of which represent potential sites of non-specific analyte binding and loss, a concern that becomes increasingly consequential as sample input amount decreases^3,9^.

A further, underappreciated complexity arises from the direction in which a trap column is operated. In forward-flush mode, the eluting mobile phase traverses the trap column in the same direction as the original sample load, meaning that any particles captured at the column inlet during loading remains permanently retained there and never reaches the analytical column; this preserves the trap column’s guard function but forces the entire eluting analyte band through the full length of the packed bed, a process expected to broaden chromatographic peaks and prolong analyte-surface contact. In reverse-flush mode, by contrast, the flow direction is reversed for elution relative to sample loading, allowing the trap column to exploit the sharp analyte focusing achieved at the inlet during loading and thereby minimize peak broadening, but at the cost of flushing any retained particles directly onto the analytical column, forfeiting the protective rationale for using a trap column in the first place. This trade-off has been noted in the proteomics literature^5,9^. A recent study systematically optimized trap column inner diameter, particle size, and loading flow rate for single-cell-level inputs, but did not vary flush direction, operating exclusively in backflush mode^11^. A side-by-side characterization spanning trap column geometry (bed length and pore size), flush direction, and sample loading conditions relevant to low-input proteomics has, to our knowledge, not yet been reported.

In contrast, direct injection removes additional surface area and dead volume from the flow path as much as possible and eliminates the risk of sample loss during a separate loading step since the sample is loaded directly onto the analytical column, often yielding the highest achievable sensitivity^1^. In a recent study, Hendriks and colleagues identified up to 7,000 proteins per single cell using direct injection, and directly benchmarked this against a trap-and-elute configuration employing a trap column; while trap-and-elute identification numbers were somewhat lower, this configuration enabled substantially higher sample throughput^12^. These findings indicate that the gap between trap-and-elute and direct injection performance is not fixed, and that it can be narrowed through careful trap column and workflow design, particularly in reverse-flush operation; however, the extent to which this gap can be closed, especially in forward-flush mode where the protective guard-column function is preserved, remains underexplored and represents an opportunity for trap column development.

In this study, we set out to systematically address this question using a purpose-built family of NanoShield™ trap column prototypes (IonOpticks) spanning multiple packed bed lengths and, ultimately, an alternative stationary phase, benchmarked against the established PepMap™ Neo trap cartridge (Thermo Fisher Scientific) and against direct injection. Using a standardized testing workflow built around a commercial Pierce™ HeLa digest standard (Thermo Fisher Scientific) and precise low-volume liquid handling using the Mosquito LV (SPT Labtech), we iteratively evaluated trap column performance across two MS platforms, multiple LC gradient lengths, both flush directions, and a range of sample input amounts and injection volumes. This iterative, feedback-driven testing process, in which performance limitations identified at each stage informed the design of the next trap column iteration, allowed us to progressively narrow the performance gap between trap-and-elute workflows and direct injection, particularly in forward-flush mode, while clarifying the underlying trade-off between analytical column protection and chromatographic performance that governs trap column design for low-input proteomics.

## RESULTS

### Detailed experimental and data-analysis methods are provided in the supplementary data

#### Overview of the testing Workflow

To enable a systematic and statistically robust comparison of trap column performance, we established a standardized testing workflow built around commercial Pierce™ HeLa digest standard^13^, precise low-volume liquid handling, and LC-MS/MS as the quantitative readout (Figure 1A). Pierce™ HeLa digest standard was used as a well-controlled, reproducible, and commercially available test sample across all experiments, avoiding batch-to-batch variability inherent to in-house sample preparation. Dilution series in sub-nanogram range for input amounts were prepared using a Mosquito low-volume liquid handler, enabling fast, accurate, and precise pipetting of sample volumes as low as tens of nanoliters. This standardized sample preparation pipeline was applied across a series of trap column benchmarking experiments comparing NanoShield™ prototypes of varying dimensions, the commercially available PepMap™ Neo trap cartridge, and direct injection, with protein and peptide identification numbers, chromatographic peak width, quantitative precision, and retention time (RT) stability as the key performance metrics. This workflow was iteratively applied across the study to interrogate trap column geometry, stationary phase chemistry, flush direction, and sample loading (input amount and injection volume), aiming to improve identification numbers and robustness while improving analytical column lifespan.

**Figure 1.**
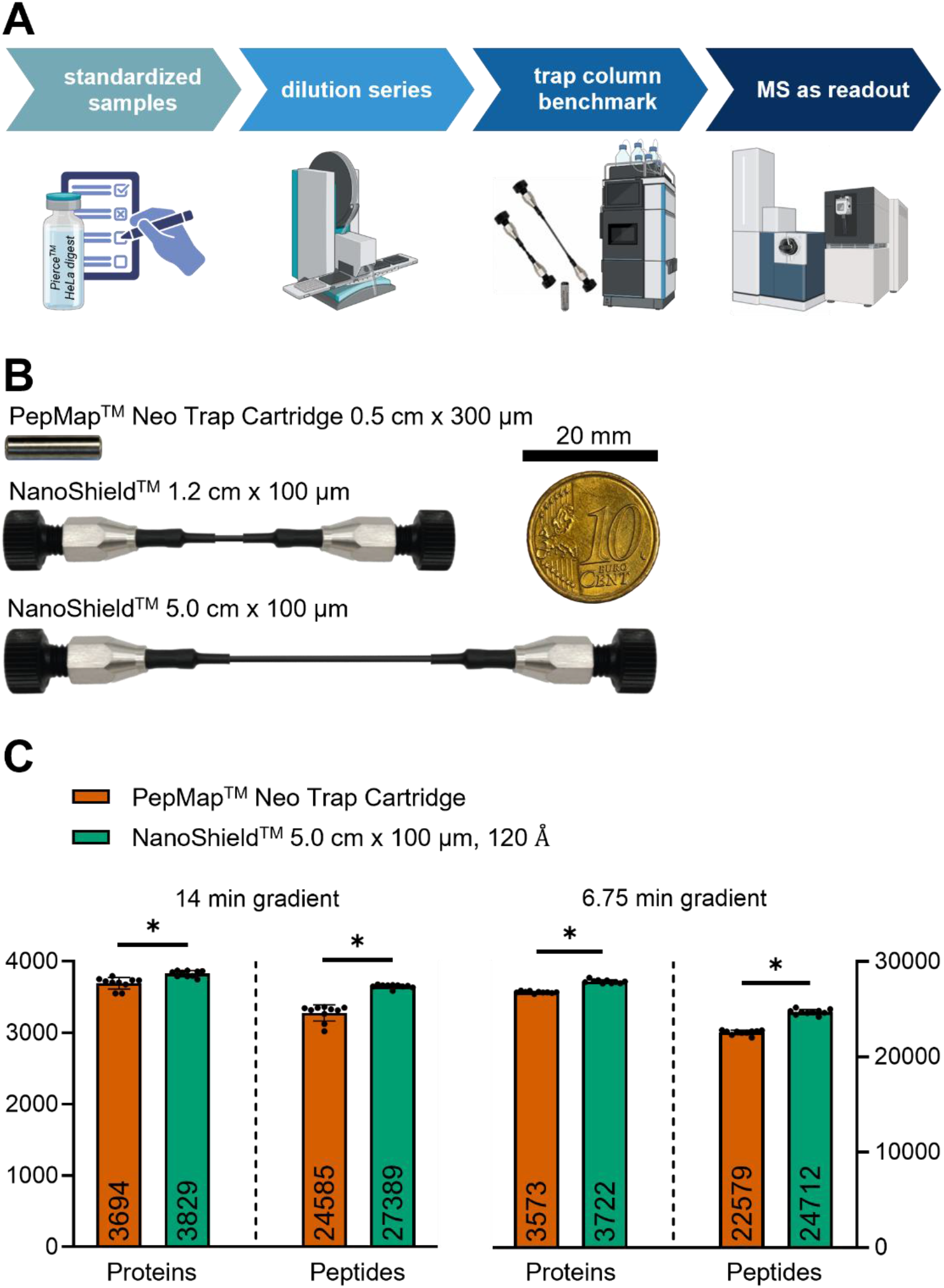
Overview of the systematic trap column testing workflow and initial benchmark (dataset a). (A) Schematic of the standardized benchmarking workflow: commercial Pierce™ HeLa digest standard was diluted to defined input amounts and injection volumes using the Mosquito LV, a low-volume liquid handler, separated by LC-MS/MS across different trap column formats, and analyzed by mass spectrometry. (B) Trap column formats evaluated in this study: PepMap™ Neo trap cartridge and NanoShield™ trap columns of two bed lengths, shown alongside a 10-cent EURO coin (diameter 19.75 mm) for size reference. Dimensions denote packed bed length × inner diameter; the trap columns themselves are considerably larger due to frits, fittings, and end-plugs. (C) Number of identified proteins (left y-axis, 1% FDR at protein level) and peptides (right y-axis, 1% FDR at precursor level) for the PepMap™ Neo trap cartridge (red) and the 5 cm NanoShield™ trap column (green), both operated in reverse-flush mode, using a 14 min or 6.75 min LC gradient with 250 pg Pierce™ HeLa digest standard. Data points represent independent sample injections (n = 10); bars and numbers show the mean; error bars show ± standard deviation (SD). Asterisks denote a statistically significant difference (p < 0.05) upon performing an unpaired t-test with Welch’s correction; significance is indicated only as present (*) or absent (ns), without implying a specific level of significance.

#### Initial benchmarking of a first-generation NanoShield™ trap column against PepMap™ Neo trap cartridge identifies a performance advantage

At the outset of this study, no short-format NanoShield™ trap column existed; the earliest available NanoShield™ prototype was a 5 cm x 100 µm column featuring a 120 Å pore size stationary phase (Figure 1B). We therefore began by comparing this NanoShield™ 5 cm x 100 µm trap column against the PepMap™ Neo trap cartridge (0.5 cm x 300 µm), a well-established trap column format, on a timsTOF Ultra 2 platform (Bruker Daltonics) (dataset a; Figure 1C). Both trap columns were operated in reverse-flush mode and tested across two LC gradient lengths, a 14 min and a shorter 6.75 min gradient, corresponding to the ∼100 and ∼30 samples per day (SPD) throughput regimes routinely used for single-cell proteomics in our laboratory, to assess whether any observed performance differences would hold across different chromatographic timescales.

Across both gradients, the NanoShield™ 5 cm trap column consistently outperformed the PepMap™ Neo trap cartridge in both protein and peptide identification numbers (Figure 1C). For the 14 min gradient, NanoShield™ yielded 3829 proteins and 27389 peptides compared to 3694 proteins and 24585 peptides for PepMap™. The same trend held for the shorter 6.75 min gradient, with NanoShield™ identifying 3722 proteins and 24712 peptides versus 3573 proteins and 22579 peptides for PepMap™. This initial, result indicated that the NanoShield™ trap column could outperform an established commercial trap column format even in its earliest, longest-format iteration, and provided the impetus to extend testing of this new trap column format to our primary analytical platform, the Orbitrap Astral (Thermo Fisher Scientific).

#### NanoShield™ outperforms PepMap™ Neo trap cartridge on the Orbitrap Astral across gradient lengths and injection volumes, but at the cost of slower sample loading

Having established a performance advantage for NanoShield™ on the timsTOF platform, we next evaluated whether this advantage would be reproduced on the Orbitrap Astral, the mass spectrometer used for all subsequent experiments in this study. We again compared the NanoShield™ 5 cm x 100 µm trap column against the PepMap™ Neo trap cartridge, both operated in reverse-flush mode across two gradient lengths (25 min and 6.25 min) (dataset b; Figure 2A, B).

**Figure 2.**
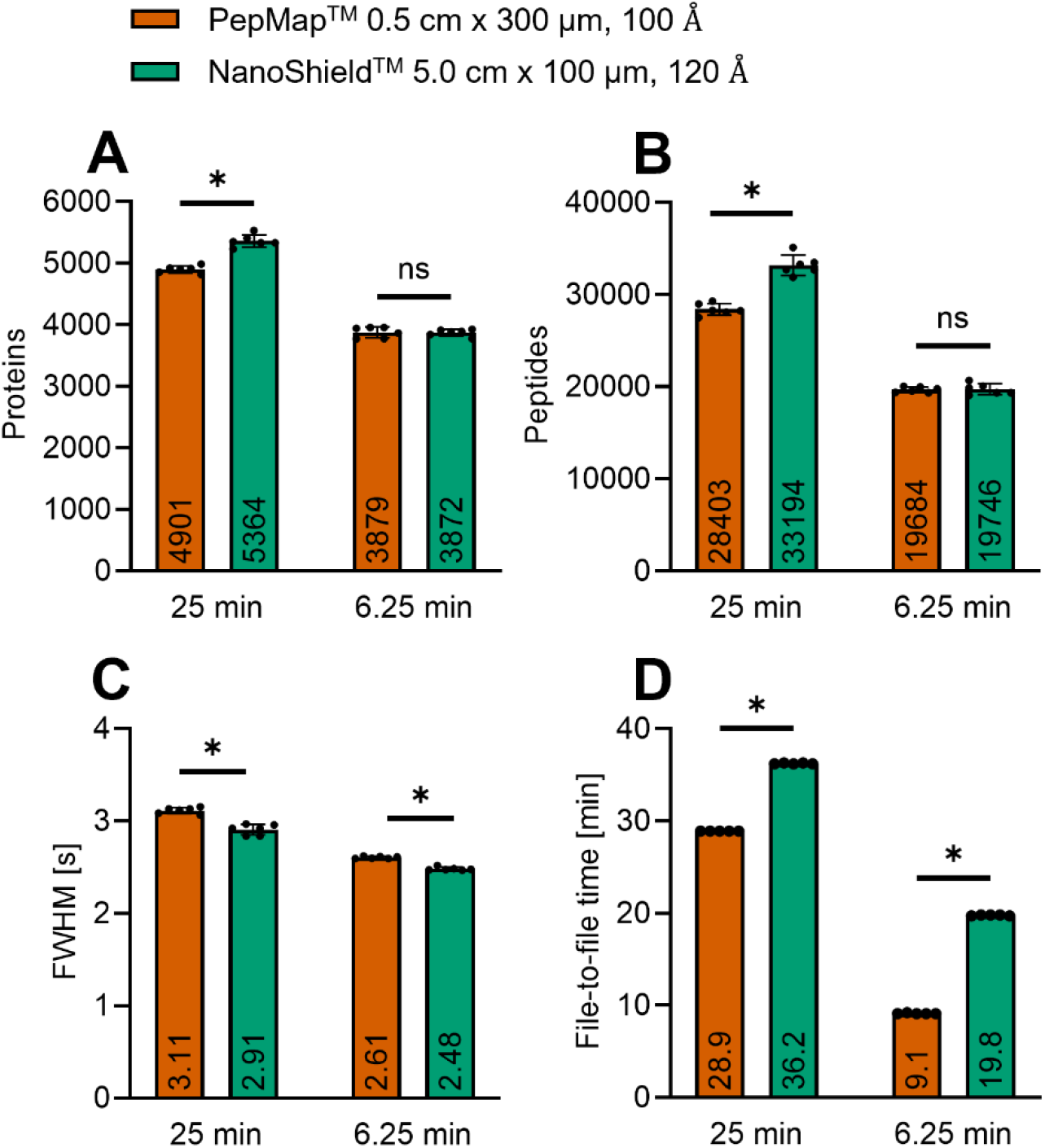
NanoShield™ 5 cm trap column outperforms PepMap™ across two gradient lengths on an Orbitrap Astral, with a trade-off in loading time (dataset b). (A, B) Number of identified proteins (A, 1% FDR at protein level) and peptides (B, 1% FDR at precursor level) for the PepMap™ Neo trap cartridge (red) and the NanoShield™ 5 cm trap column (green), both operated in reverse-flush mode, using a 25 min (left) or 6.25 min (right) LC gradient; legend dimensions denote packed bed length x inner diameter, stationary phase pore size. (C) Average chromatographic peak width (full width half maximum, FWHM) for the corresponding runs. (D) File-to-file time, defined as the total elapsed time between successive raw file writes, encompassing sample loading, equilibration, separation, and column wash. 250 pg Pierce™ HeLa digest standard per injection. Data points represent independent sample injections (n = 6, for file-to-file time: n = 5); bars and numbers show the mean; error bars show ± SD. Asterisks denote a statistically significant difference (p < 0.05) upon performing an unpaired t-test with Welch’s correction; significance is indicated only as present (*) or absent (ns), without implying a specific level of significance.

This experiment recapitulated the trend observed on the timsTOF platform: for the longer, 25 min gradient, NanoShield™ outperformed PepMap™ (5364 versus 4901 proteins). Notably, this performance gap largely vanished for the shorter, 6.25 min gradient, where both trap columns performed essentially equivalently (3872 versus 3879 proteins), suggesting that at short gradient times, elution speed becomes limiting for both trap columns alike, masking any trap column specific differences that are otherwise apparent at longer gradients. Average chromatographic peak width (FWHM) was comparable or marginally narrower for NanoShield™ at the 25 min gradient (2.91 versus 3.11 s), with no clear difference at the 6.25 min gradient (Figure 2C), providing additional, if modest, evidence of superior LC performance for the NanoShield™ format at this stage.

However, this performance benefit came with a practical cost: file-to-file time, the total elapsed time between successive raw file writes encompassing sample loading, equilibration, separation, and column wash, was substantially longer for the NanoShield™ than for the PepMap™ across both gradients (e.g., 36.2 versus 28.9 min for the 25 min gradient; Figure 2D). Isolated backpressure measurements confirmed that this originated from the substantially higher backpressure generated by the 5 cm NanoShield™ trap column during sample loading compared to the PepMap™ Neo trap cartridge (Supplementary Figure 2). This, at a constant controlled loading pressure of 800 bar (IonOpticks recommended loading pressure, applied uniformly across all trap column experiments for fair comparison), necessarily reduces the achievable loading flow rate and thus extends sample loading time. Because sample loading time scales directly with trap column backpressure at fixed loading pressure, and because a longer packed bed carries an inherently higher risk of irreversible sample loss through non-specific binding, we reasoned that a shorter NanoShield™ trap column of identical inner diameter and stationary phase pore size should reduce backpressure and loading time while preserving the diameter-dependent advantages already observed. A shorter trap column bed was further expected to reduce the peak broadening associated with operating a trap column in forward-flush mode, since forward-flush elution forces the full sample volume through the entire length of the packed bed, an effect that should scale with trap column length. These considerations motivated us to next evaluate a substantially shortened NanoShield™ trap column format (Figure 2B), and to explicitly benchmark trap column performance under both forward- and reverse-flush operation, a comparison that would prove central to the remainder of this study.

#### A shortened NanoShield™ format closes the loading time gap, improves forward-flush performance, and approaches direct injection level performance

To test the hypothesis that reducing trap column length would mitigate the loading time penalty and forward-flush peak broadening observed for the 5 cm NanoShield™ format, we performed a back-to-back comparison of three trap columns, PepMap™ Neo trap cartridge, the original NanoShield™ 5 cm x 100 µm and a newly available, substantially shortened NanoShield™ 1.2 cm x 100 µm, each operated in both forward- and reverse-flush mode on an Orbitrap Astral (dataset c; Figure 3A–D).

**Figure 3.**
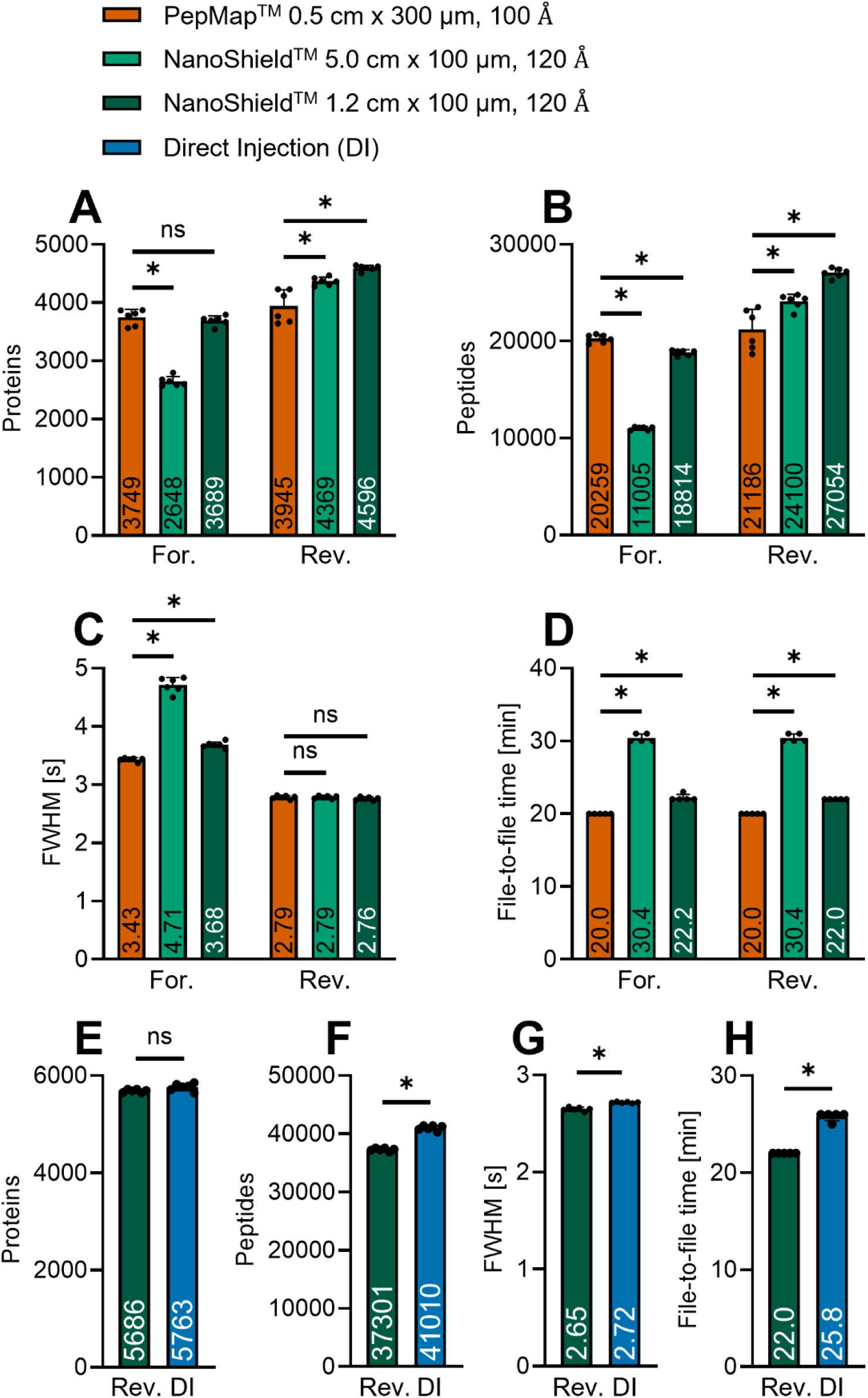
Shortening the NanoShield™ trap column narrows the forward-flush performance gap, and reverse-flush operation approaches direct injection performance (datasets c, d). (A–D) Number of identified proteins (A, 1% FDR at protein level), peptides (B, 1% FDR at precursor level), average chromatographic peak width (C, FWHM), and file-to-file time (D) for the PepMap™ Neo trap cartridge (red), the NanoShield™ 5 cm trap column (light green), and the NanoShield™ 1.2 cm trap column (dark green) operated in forward- and reverse-flush mode (For., Rev.), using a 14 min LC gradient; legend dimensions denote packed bed length x inner diameter, stationary phase pore size (dataset c). (E–H) Number of identified proteins (E), peptides (F), average chromatographic peak width (G, FWHM), and file-to-file time (H) for the NanoShield™ 1.2 cm trap column operated in reverse-flush mode (dark green) and direct injection (DI, blue), using the same 14 min LC gradient (dataset d). 250 pg Pierce™ HeLa digest standard per injection. Data points represent independent sample injections (n = 6); bars and numbers show the mean; error bars show ± SD. Asterisks denote a statistically significant difference (p < 0.05) upon performing an unpaired t-test with Welch’s correction; significance is indicated only as present (*) or absent (ns), without implying a specific level of significance.

In reverse-flush mode, both NanoShield™ formats outperformed PepMap™ in protein and peptide identifications, with the shortened 1.2 cm NanoShield™ yielding the highest numbers overall (4596 proteins, 27054 peptides), ahead of the 5 cm NanoShield™ (4369 proteins, 24100 peptides) and PepMap™ (3945 proteins, 21186 peptides; Figure 3A, B). In forward-flush mode, however, the 5 cm NanoShield™ suffered a pronounced drop in performance (2648 proteins, 11005 peptides), falling well below PepMap™ (3749 proteins, 20259 peptides), consistent with substantial peak broadening in this longer trap format, as directly evidenced by its markedly increased FWHM (4.71 s versus 3.43 s for PepMap™; Figure 3C). The shortened 1.2 cm NanoShield™, in contrast, largely recovered forward-flush performance, statistically indistinguishable from PepMap™ in identification numbers (3689 proteins, 18814 peptides) and producing a correspondingly much narrower FWHM (3.68 s) than its 5 cm predecessor. The 5 cm NanoShield™ had already outperformed PepMap™ in reverse-flush mode, and the shortened 1.2 cm NanoShield™ further outperformed the 5 cm format in reverse-flush mode while matching PepMap™ in forward-flush mode. The 1.2 cm NanoShield™ therefore emerged as the best-performing trap column overall across both flush directions tested in this experiment.

Reasoned by reduced backpressure and therefore faster achievable loading flow rate at the same controlled loading pressure of 800 bar, the shortened NanoShield™ showed a substantial reduction in file-to-file time relative to the 5 cm format (22.2 versus 30.4 min in forward-flush mode; 22.0 versus 30.4 min in reverse-flush mode; Figure 3D, Supplementary Figure 2). As expected, file-to-file time was unaffected by flush direction for any trap column format, since sample loading, the rate-limiting step, is identical regardless of the subsequent elution direction. Taken together, these results demonstrate that shortening the NanoShield™ trap column from 5 cm to 1.2 cm, while retaining the same 100 µm inner diameter and 120 Å pore size of the stationary phase, simultaneously improves forward-flush performance, reduces sample loading time, and retains or improves upon the reverse-flush identification advantage over the PepMap™ Neo trap cartridge, establishing the shortened NanoShield™ as the leading trap column candidate for all subsequent experiments.

Having identified the 1.2 cm NanoShield™ operated in reverse-flush mode as the best-performing trap-and-elute configuration tested thus far, we next asked how this configuration compares to direct injection, generally regarded as the most sensitive sample introduction strategy for low-input proteomics since it avoids any trap column-associated dead volume or surface-binding losses. We therefore performed a back-to-back comparison of the NanoShield™ 1.2 cm x 100 µm trap column, operated exclusively in reverse-flush mode, against direct injection on the Orbitrap Astral (dataset d; Figure 3E–H).

NanoShield™ in reverse-flush mode closely approached direct injection performance across all metrics tested. Protein identification numbers were statistically indistinguishable between the two conditions (5686 versus 5763 proteins; Figure 3E), while peptide numbers remained modestly higher for direct injection (37301 versus 41010 peptides; Figure 3F). Chromatographic peak width (FWHM) was comparably narrow for both conditions (2.65 versus 2.72 s; Figure 3G), indicating that the trap column introduced no detectable additional band broadening relative to direct injection. Notably, file-to-file time was shorter for NanoShield™ than for direct injection (22.0 versus 25.8 min; Figure 3H), reflecting the faster achievable sample loading for the given low sample volumes. Together, these results show that, at this stage of trap column optimization, the shortened NanoShield™ in reverse-flush mode had essentially closed the performance gap to direct injection while retaining a throughput advantage, establishing it as a compelling trap-and-elute alternative to direct injection for low-input proteomic workflows.

#### NanoShield™ performance advantage is robust across sample input amount and injection volume

With the shortened NanoShield™ trap column established as a strong, direct injection competitive performer under standard loading conditions, we next assessed how robust this performance advantage is to the two sample loading parameters most relevant to practical low-input proteomics workflows: the absolute amount of sample loaded and the volume in which that sample is delivered. Low-input samples are frequently constrained in total peptide input, and the required injection volume can vary considerably; a trap column intended for routine low-input use must tolerate this variation in both parameters without a disproportionate loss in performance. We addressed these two parameters in two separate titration experiments on the Orbitrap Astral, comparing the PepMap™ Neo trap cartridge, the shortened NanoShield™ and direct injection. A 1.5 cm x 100 µm prototype with 120 Å pore size functionally equivalent to the 1.2 cm format was used. The small difference in packed bed length reflects prototype-to-prototype manufacturing variability rather than a deliberate design change. For this test, trap-and-elute columns were operated in reverse-flush mode.

In the first experiment, we titrated sample input amount across a 25–250 pg range at a constant injection volume of 1 µL (dataset e; Supplementary Figure 1A–D). NanoShield™ consistently outperformed PepMap™ in protein and peptide identifications across the entire input range tested (e.g., proteins at 25 pg: 3067 versus 2644; at 250 pg: 4572 versus 4027), with direct injection yielding the highest identification numbers across the full range of input amounts tested. For peptides, the gap between trap-and-elute to direct injection was considerably more pronounced than for proteins: at 25 pg, direct injection yielded 24806 peptides compared to 14260 for NanoShield™ and 11258 for PepMap™, a gap that narrowed but remained substantial at higher input amounts (e.g., at 250 pg: 30900 for direct injection versus 24023 for NanoShield™ and 19322 for PepMap™), indicating that trap-and-elute workflows disproportionately under-identify peptides relative to proteins compared to direct injection at low sample input. Chromatographic peak width (FWHM) was essentially stable across the input range for all three conditions, with direct injection producing the narrowest peaks, NanoShield™ the widest, and PepMap™ intermediate but only marginally different from NanoShield™. The one metric for which PepMap™ had a clear lead over NanoShield™ was file-to-file time; PepMap™ remained fastest across the entire titration, followed by NanoShield™, with direct injection consistently slowest, mirroring the loading time trade-off described above.

In the second experiment, we titrated sample injection volume across a 50–4000 nL range at a constant sample input amount of 250 pg (dataset f; Supplementary Figure 1E–H). NanoShield™ again outperformed PepMap™ in protein identifications across the full sample volume range tested (e.g., at 50 nL: 3859 versus 3063; at 4000 nL: 3730 versus 3189) and notably exceeded even direct injection at the lower injection volumes (50–1000 nL), with direct injection only overtaking NanoShield™ at the highest injection volume tested (4000 nL: 3964 versus 3730 proteins). Peptide numbers followed a similar pattern, with NanoShield™ again ahead of PepMap™ throughout (e.g., at 50 nL: 15932 versus 13967 peptides), while direct injection yielded comparable or higher peptide numbers at most injection volumes tested, most notably at 4000 nL. Like the input titration, chromatographic peak width (FWHM) was broadest for NanoShield™ across the entire volume range (3.14–3.38 sec), markedly wider than for PepMap™, which remained narrowest and essentially constant (2.74 s), and for direct injection, which was intermediate (3.00–3.02 s). For file-to-file time, NanoShield™ again occupied an intermediate position, slower than PepMap™ but consistently faster than direct injection across the volume range, reflecting the expected ordering of trap column backpressure and direct injection loading constraints. Together, these two titration series demonstrate that the performance advantage of the shortened NanoShield™ trap column over PepMap™ Neo trap cartridge is maintained robustly across a wide range of both sample input amounts and injection volumes, underscoring its suitability for the variable loading conditions typical of low-input proteomic sample preparation.

#### A next-generation NanoShield™ trap column with a new stationary phase narrows the forward-flush performance gap and matches direct injection level performance in reverse-flush mode

Across every trap column comparison presented thus far, reverse-flush operation consistently outperformed forward-flush operation, most pronounced for the original 5 cm NanoShield™ format (Figure 3A–D) and evident, to a lesser extent, for the shortened NanoShield™ and PepMap™ Neo trap cartridge. Across every trap column comparison presented thus far, reverse-flush operation consistently outperformed forward-flush operation, most pronounced for the original 5 cm NanoShield™ format (Figure 3A–D) and evident, to a lesser extent, for the shortened NanoShield™ and PepMap™ Neo trap cartridge. Subsequently, a next-generation NanoShield™ stationary phase was developed to improve forward-flush performance. This new stationary phase features a substantially larger pore size (300 Å, compared to 120 Å for the original NanoShield™ stationary phase and 100 Å for the PepMap™ Neo trap cartridge; Supplementary Table 1) while retaining the same particle size, inner diameter, and pressure rating as the previous NanoShield™ generation. We designed a final testing series, using a 14 min LC gradient, to evaluate the extent to which this larger pore size stationary phase narrows the forward-versus reverse-flush performance gap, and to benchmark this next-generation NanoShield™ 1.2 cm x 100 µm, 300 Å trap column against the PepMap™ Neo trap cartridge and direct injection (dataset g; Figure 4).

**Figure 4.**
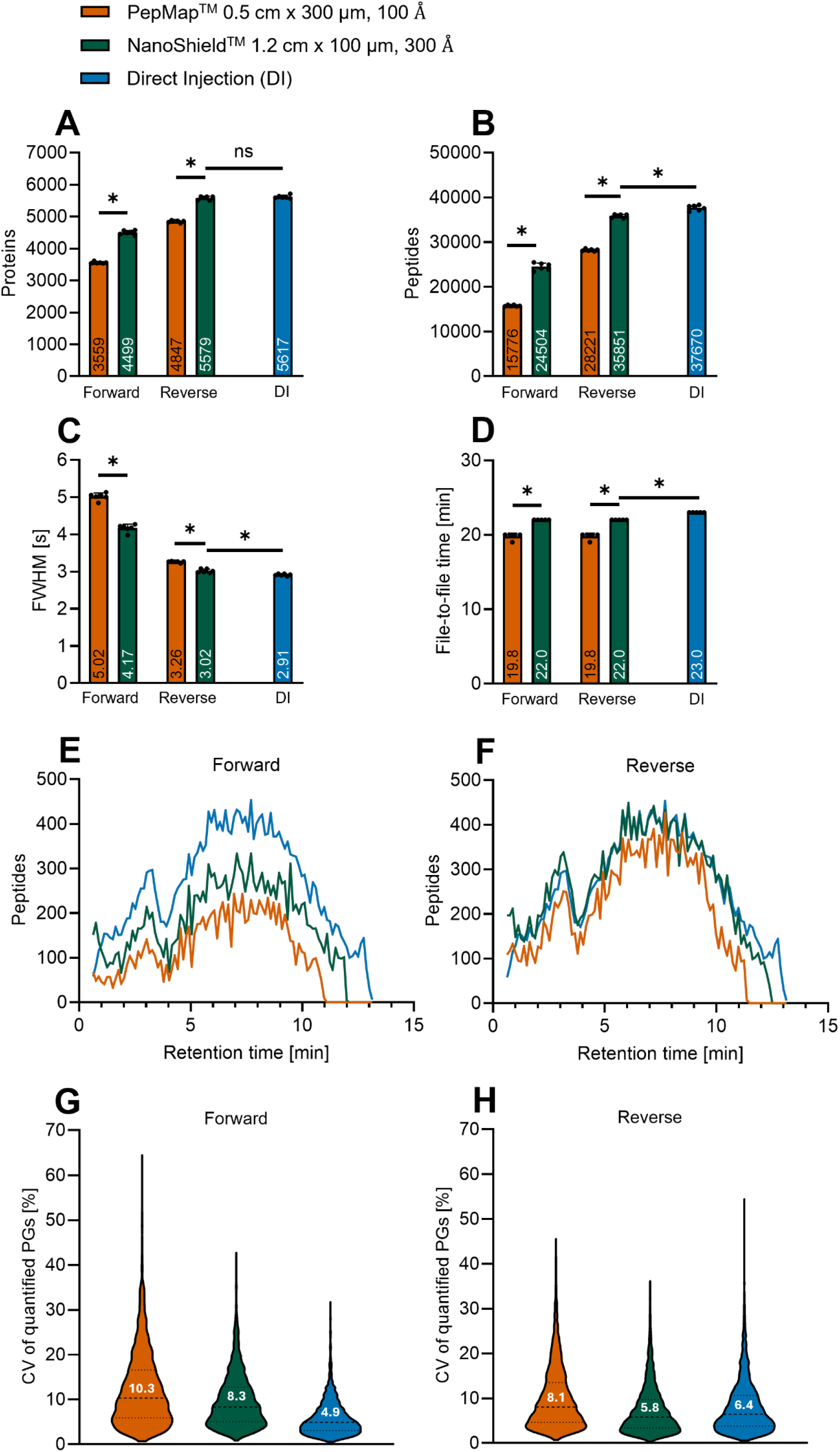
A next-generation NanoShield™ stationary phase improves, but does not fully equalize, forward-flush performance relative to reverse-flush mode (dataset g). (A, B) Number of identified proteins (A, 1% FDR at protein level) and peptides (B, 1% FDR at precursor level) for the PepMap™ Neo trap cartridge (red) and the NanoShield™ 1.2 cm trap column with the new stationary phase (green), and direct injection (DI, blue), operated in forward- and reverse-flush mode, using a 14 min LC gradient; legend dimensions denote packed bed length x inner diameter, stationary phase pore size. (C) Average chromatographic peak width (FWHM). (D) File-to-file time, (E, F) number of identified peptides across the LC gradient in forward-(E) and reverse-flush (F) mode, binned by RT (100 bins); peptides were required to be identified in every replicate of a condition, and trap-column RT were aligned to the direct injection scale using an isotonic calibration on shared anchor peptides. (G, H) Distribution of coefficients of variation (CV) of quantified protein groups (PGs) across replicate injections for each condition in forward-flush (G) and reverse-flush (H) mode, shown as violin plots with the median CV indicated inside each violin. 250 pg Pierce™ HeLa digest standard per injection. Data points represent independent sample injections (n = 6); bars and numbers show the mean; error bars show ± SD. Asterisks denote a statistically significant difference (p < 0.05) upon performing an unpaired t-test with Welch’s correction; significance is indicated only as present (*) or absent (ns), without implying a specific level of significance.

This new NanoShield™ trap column has a manufacturer-specified preferred flow direction, communicated by IonOpticks, although the column remains operable in either orientation. In preliminary testing we found that mounting the trap column so that its preferred direction aligned with forward-flush elution improved forward-flush performance without compromising reverse-flush performance; we therefore adopted this single, forward-preferred orientation for both the forward- and reverse-flush measurements presented here, simplifying the resulting comparison.

Across both protein and peptide identification numbers, the NanoShield™ 300 Å trap column outperformed the PepMap™ Neo trap cartridge in both forward- and reverse-flush mode (Figure 4A, B). In forward-flush mode, NanoShield™ (4499 proteins, 24504 peptides) exceeded PepMap™ (3559 proteins, 15776 peptides). In reverse-flush mode, NanoShield™ (5579 proteins, 35851 peptides) again exceeded PepMap™ (4847 proteins, 28221 peptides), and approached direct injection (5617 proteins, 37670 peptides), with no significant difference between NanoShield™ in reverse-flush mode and direct injection in protein identification numbers. Despite this improvement in forward-flush performance relative to the previous NanoShield™ generation, reverse-flush operation remained superior to forward-flush operation for every trap column format tested in this experiment, including the new NanoShield™ 300 Å variant (4499 versus 5579 proteins, forward versus reverse; Figure 4A), confirming that while the larger pore size stationary phase mitigates the forward-flush performance penalty, it does not eliminate it.

This residual penalty in performance is consistent with multiple compounding mechanisms inherent to forward-flush elution: the full sample volume is forced through the entire packed bed length, exacerbating peak broadening; the extended transit time and surface contact with the full trap column bed increases the opportunity for peptide loss through non-specific binding; and the focusing effect achieved during sample loading at the trap column inlet, which is efficiently preserved and exploited during reverse-flush elution, but is substantially diminished when the sample is instead eluted forward through the trap column bed.

Chromatographic peak width (FWHM) mirrored this pattern: the NanoShield™ 300 Å trap column produced narrower peaks than the PepMap™ Neo trap cartridge in both flush directions (forward: 4.17 versus 5.02 s; reverse: 3.02 versus 3.26 s), and in reverse-flush mode achieved FWHM values nearly indistinguishable from direct injection (3.02 versus 2.91 s; Figure 4C). Forward-flush mode produced substantially broader peaks for every trap column tested, reflecting the peak-broadening mechanism described above. File-to-file time was unaffected by flush direction for any condition, as expected given that sample loading, rather than elution direction, is rate-limiting (Figure 4D); PepMap™ remained faster than the NanoShield™ 300 Å trap column (19.8 versus 22.0 min in both flush directions), consistent with its lower measured backpressure (Supplementary Figure 2), while the NanoShield™ 300 Å trap column in turn remained faster than direct injection (23.0 min).

Resolving peptide identification numbers across the RT axis provided a complementary view of this same identification advantage (Figure 4E, F). All conditions followed the same overall elution profile, dictated by the analytical column, gradient, and sample, but direct injection identified more peptides than either trap column at essentially every point across the RT range, with the NanoShield™ 300 Å trap column consistently identifying more peptides than the PepMap™ Neo trap cartridge throughout the full gradient. The gap between direct injection and the two trap columns was more pronounced in forward-flush mode (Figure 4E) than in reverse-flush mode, where the three profiles converged much more closely (Figure 4F), further corroborating the greater forward-flush performance penalty for trap-and-elute workflows described above.

Quantitative precisions, assessed via the distribution of coefficient of variation (CV) of protein groups quantified further reinforced this picture: in forward-flush mode, PepMap™ showed the broadest and highest median CV distribution (10.3%), the NanoShield™ 300 Å trap column showed a tighter distribution (8.3%), and direct injection showed the tightest (4.9%; Figure 4G). In reverse-flush mode, this gap between trap columns and direct injection narrowed considerably, with the NanoShield™ 300 Å trap column achieving the lowest median CV of all conditions tested (5.8%, compared to 6.4% for direct injection and 8.1% for PepMap™; Figure 4H), indicating excellent quantitative precision for the NanoShield™ 300 Å trap column under its optimal operating configuration.

RT stability across replicate injections was assessed as the per-precursor SD of RT, restricted to precursors identified in every replicate of every condition tested (Supplementary Figure 3A, B). In forward-flush mode, RT variability differed meaningfully between conditions (Friedman test, p < 0.001, Kendall’s W = 0.353): PepMap™ showed the broadest spread (median SD 0.0199 min), while the NanoShield™ 300 Å (median SD 0.0127 min) was more stable and approached the direct injection reference (median SD 0.0112 min), with all pairwise comparisons statistically significant. In reverse-flush mode, all three conditions produced tight, comparable distributions (medians 0.0094–0.0113 min); despite being statistically significant, the associated effect size was negligible (Kendall’s W = 0.039), indicating a negligible difference. These results show a genuine, moderate improvement in RT stability for the short, large pore size NanoShield™ trap column over PepMap™ Neo trap cartridge in forward-flush mode, while in reverse-flush mode RT stability is essentially indistinguishable between trap columns and direct injection.

Taken together, this final dataset demonstrates that the next-generation NanoShield™ trap column, featuring a new stationary phase with increased pore size (from 120 Å to 300 Å) and developed in response to the forward-flush performance penalty identified earlier, provides a consistent performance advantage over the PepMap™ Neo trap cartridge across identification numbers, peak width, and quantitative precision in both flush directions, and achieves a measurable, though incomplete, narrowing of the forward-versus reverse-flush performance gap. In its optimal configuration, reverse-flush mode, the NanoShield™ 300 Å trap column matches or exceeds direct injection level performance in peak width and quantitative precision while retaining markedly higher protection potential, underscoring that iterative, feedback-driven trap column engineering can progressively close the performance gap between trap-and-elute workflows and the current direct injection gold standard for low-input proteomics.

## DISCUSSION

This study presents a systematic, chronologically structured benchmarking effort that tracked the iterative optimization of a new trap column format, the NanoShield™ trap column from IonOpticks. Starting from an initial 5 cm prototype, we progressed through increasingly shorter, large pore size geometries in direct response to performance limitations identified at each stage. The resulting shortened NanoShield™ trap column with a stationary phase featuring a pore size of 300 Å operated in reverse-flush mode, approached or matched direct injection performance across identification numbers, peak width, and quantitative precision. Below, we discuss the mechanistic and practical implications of this trajectory and its current limitations.

### Trap column length and flush direction jointly govern the protection vs performance trade-off

The consistent superiority of reverse- over forward-flush operation, observed across every trap column format tested, is readily rationalized by the fluid dynamics of trap-and-elute chromatography: reverse-flush exploits the analyte focusing achieved at the trap column inlet during sample loading, whereas forward-flush forces the entire analyte band through the full packed bed, compounding peak broadening and prolonging analyte-surface contact time and thus non-specific binding losses^5,8–10^. That shortening the NanoShield™ bed from 5 cm to 1.2 cm essentially eliminated the forward-flush identification deficit relative to the much shorter PepMap™ Neo trap cartridge indicates that bed length is a primary determinant of forward-flush performance. The data acquired for the short NanoShield™ with a stationary phase featuring a larger pore size of 300 Å add a second, partially independent lever: pore size further narrowed, but did not eliminate, the forward-flush penalty, suggesting a mechanism rooted in intra-particle mass transfer and surface interactions^14,15^, distinct from the band-broadening effect of bed length itself. Taken together, these findings already offer actionable, present-day guidance: where guard-column protection is not required, a short trap column operated in reverse-flush mode is the better-performing default; where sample matrix complexity necessitates forward-flush protection, minimizing trap column length and favoring larger pore size stationary phases are the most effective available levers for limiting the associated performance cost. However, bed length also sets loading capacity, so any reduction in trap column length must be balanced against the sample input amounts the trap is expected to accommodate, see Supplementary Table 1 for trap column specific loading capacities.

### A shortened, large pore size trap column narrows the gap between trap-and-elute and direct injection

A central motivation for this study was to determine whether trap-and-elute workflows could approach the sensitivity of direct injection^1,12^ without forfeiting analytical column protection. Our data show that this is largely achievable in reverse-flush mode, where the optimized NanoShield™ trap column became nearly indistinguishable from direct injection in peak width and quantitative precision, and competitive in identification numbers, while retaining a throughput advantage from faster sample loading. This performance parity, however, is achieved precisely in the operating mode that forfeits full guard column function; the choice between forward- and reverse-flush therefore remains a deliberate trade-off between sensitivity and analytical column protection, rather than one resolved by trap column design alone. Robustness across a wide range of sample input amounts and injection volumes further supports that this advantage is not contingent on idealized loading conditions, strengthening its relevance for routine, variable low-input workflows.

### Limitations and outlook

Several limitations temper these conclusions and define clear next steps. Most importantly, this study did not assess long-term robustness: we evaluated trap column performance over single to few injections, but not the extent to which forward-flush operation extends analytical column lifespan under routine, repeated use, nor how trap column and direct injection performance evolve over hundreds of injections, though guard-column protection is established practice for extending column lifespan in LC more broadly.^8,16^ Additionally, nearly all experiments were conducted at single-cell-level sample loads; at higher sample inputs, the trade-off between forward- and reverse-flush operation is likely less pronounced. Establishing this would allow a more quantitative answer to when forward-flush protection is truly necessary, and when a trap column’s performance cost is justified at all relative to direct injection. Second, all experiments used a single, well-characterized HeLa digest standard at low-input amounts; performance, particularly non-specific binding losses and the forward-flush penalty, may differ substantially for more complex or higher-input sample matrices, and has not yet been tested for multiplexed or crosslinked proteomics workflows, where sample complexity and dynamic range differ markedly from the conditions evaluated here. Third, quantitative accuracy was not directly benchmarked across PepMap™, NanoShield™, and direct injection using defined proteome mixes, so while our comparisons capture identification numbers, peak width, and quantitative precision, they do not address whether quantitative accuracy differs systematically between trap configurations and direct injection — a consideration that may be particularly relevant for quantitative experiments such as differential expression studies. Addressing these points, particularly long-term lifespan testing and extension to higher input and more complex sample types, represents the logical next step of this work and would allow the present, still largely qualitative design guidance to be translated into quantitative, application-specific recommendations for trap column choice and operating mode.

In conclusion, this work demonstrates that iterative, feedback-driven trap column engineering can progressively narrow the trade-off between analytical column protection and chromatographic performance in trap-and-elute workflows, with trap column length and stationary phase pore size emerging as key, partially independent design levers. Long-term robustness, performance under higher-input and more complex sample types, and quantitative accuracy relative to direct injection remain to be established, representing clear next steps as the single-cell and low-input proteomics field continues to move toward increasingly diverse, lower-input sample types^17,18^.

## SUPPLEMENTARY DATA

This article contains supplementary data

## ACKNOWLEDGEMENTS

During the preparation of this work, the authors used Claude Sonnet 5 available in Abacus (abacus.ai) to draft initial versions of the manuscript. The authors substantially revised the generated text, verified all factual content against the source material, and take full responsibility for the content of the publication.

## AUTHOR CONTRIBUTIONS

D.D. and M.M. conceptualized the study. D.D. performed experiments, analyzed the data, and wrote the manuscript. L.C. designed the NanoShield™ trap columns, and J.J.S. and T.B. oversaw their development. T.B. and L.C. supplied the prototypes and provided feedback on the design and experiments of the present study. M.M. and K.M. supervised the study. All authors revised, improved, and agreed on the manuscript.

## ABBREVIATIONS

CV: Coefficient of Variation
DI: Direct Injection
FDR: False Discovery Rate
FWHM: Full Width at Half Maximum
LC: Liquid Chromatography
LC-MS: Liquid Chromatography–Mass Spectrometry
MS: Mass Spectrometry
RT: Retention Time
SCP: Single-Cell Proteomics
SD: Standard Deviation
SPD: Samples Per Day

## REFERENCES

1. Bubis, J. A. et al. Challenging the Astral mass analyzer to quantify up to 5,300 proteins per single cell at unseen accuracy to uncover cellular heterogeneity. Nat. Methods 22, 510–519 (2025).

2. Karagach, S., Smollich, J., Atrakchi, O., Mohan, V. & Geiger, T. High-Throughput Single-Cell Proteomics of In Vivo Cells. Mol. Cell. Proteomics 24, 101018 (2025).

3. Petrosius, V. et al. Exploration of cell state heterogeneity using single-cell proteomics through sensitivity-tailored data-independent acquisition. Nat. Commun. 14, 5910 (2023).

4. Leodolter, J., Thierer, T., Mechtler, K. & Matzinger, M. Pushing the limits of single-cell proteomics to investigate bacterial heterogeneity using bacSCP. Nat. Commun. 17, 10425 (2026).

5. Zheng, R. et al. A High-Sensitivity Low-Nanoflow LC-MS Configuration for High-Throughput Sample-Limited Proteomics. Anal. Chem. 95, 18673–18678 (2023).

6. Bian, Y. et al. Robust, reproducible and quantitative analysis of thousands of proteomes by micro-flow LC–MS/MS. Nat. Commun. 11, 157 (2020).

7. Shen, Y. et al. High-efficiency nanoscale liquid chromatography coupled on-line with mass spectrometry using nanoelectrospray ionization for proteomics. Anal. Chem. 74, 4235–4249 (2002).

8. Miletić Vukajlović, J., Ilić, B., Bruszel, B., Panić-Janković, T. & Mitulović, G. Comparison of the Trapping Efficiency for Tryptic Peptides on Particle-Packed and Micro-Pillar Trap Columns for Proteomics Analyses. Proteomes 14, 10 (2026).

9. Webber, K. G. I. et al. Open-tubular trap columns: towards simple and robust liquid chromatography separations for single-cell proteomics. *Mol*. Omics 20, 184–191 (2024).

10. Wanninger, M. Columns Matter, Too: How to Improve Nano and Microflow LC-MS Proteomics by Considering the Separation Itself - Waters Blog. https://www.waters.com/blog/ https://www.waters.com/blog/columns-matter-too-how-to-improve-nano-and-microflow-lc-ms-proteomics-by-considering-the-separation-itself/ (2018).

11. Huang, S. et al. Optimization of trap column properties and loading conditions for proteome profiling of single-cell-level sample inputs. Anal. Bioanal. Chem. 418, 4801–4810 (2026).

12. Hendriks, I. A. et al. Extending single-cell proteomics from HeLa to small primary human immune cells. Nat. Commun. 17, 10985 (2026).

13. Köcher, T., Pichler, P., Swart, R. & Mechtler, K. Analysis of protein mixtures from whole-cell extracts by single-run nanoLC-MS/MS using ultralong gradients. Nat. Protoc. 7, 882–890 (2012).

14. Wei, B., Zhang, B., Boyes, B. & Zhang, Y. T. Reversed-phase chromatography with large pore superficially porous particles for high throughput immunoglobulin G2 disulfide isoform separation. J. Chromatogr. A 1526, 104–111 (2017).

15. Bobály, B., Veuthey, J.-L., Guillarme, D. & Fekete, S. New developments and possibilities of wide-pore superficially porous particle technology applied for the liquid chromatographic analysis of therapeutic proteins. J. Pharm. Biomed. Anal. 158, 225–235 (2018).

16. Dolan, J. W. Column Protection, Three Easy Steps | LCGC International. https://www.chromatographyonline.com/view/column-protection-three-easy-steps-1 (2026).

17. Momenzadeh, A. & Meyer, J. G. Single-cell proteomics using mass spectrometry. Cell Genomics 5, 100973 (2025).

18. Sanchez-Avila, X., de Oliveira, R. M., Huang, S., Wang, C. & Kelly, R. T. Trends in Mass Spectrometry-Based Single-Cell Proteomics. Anal. Chem. 97, 5893–5907 (2025).

