## Supplementary for "Trapping more seeing more, benchmarking a new trap column for low-input proteomics"

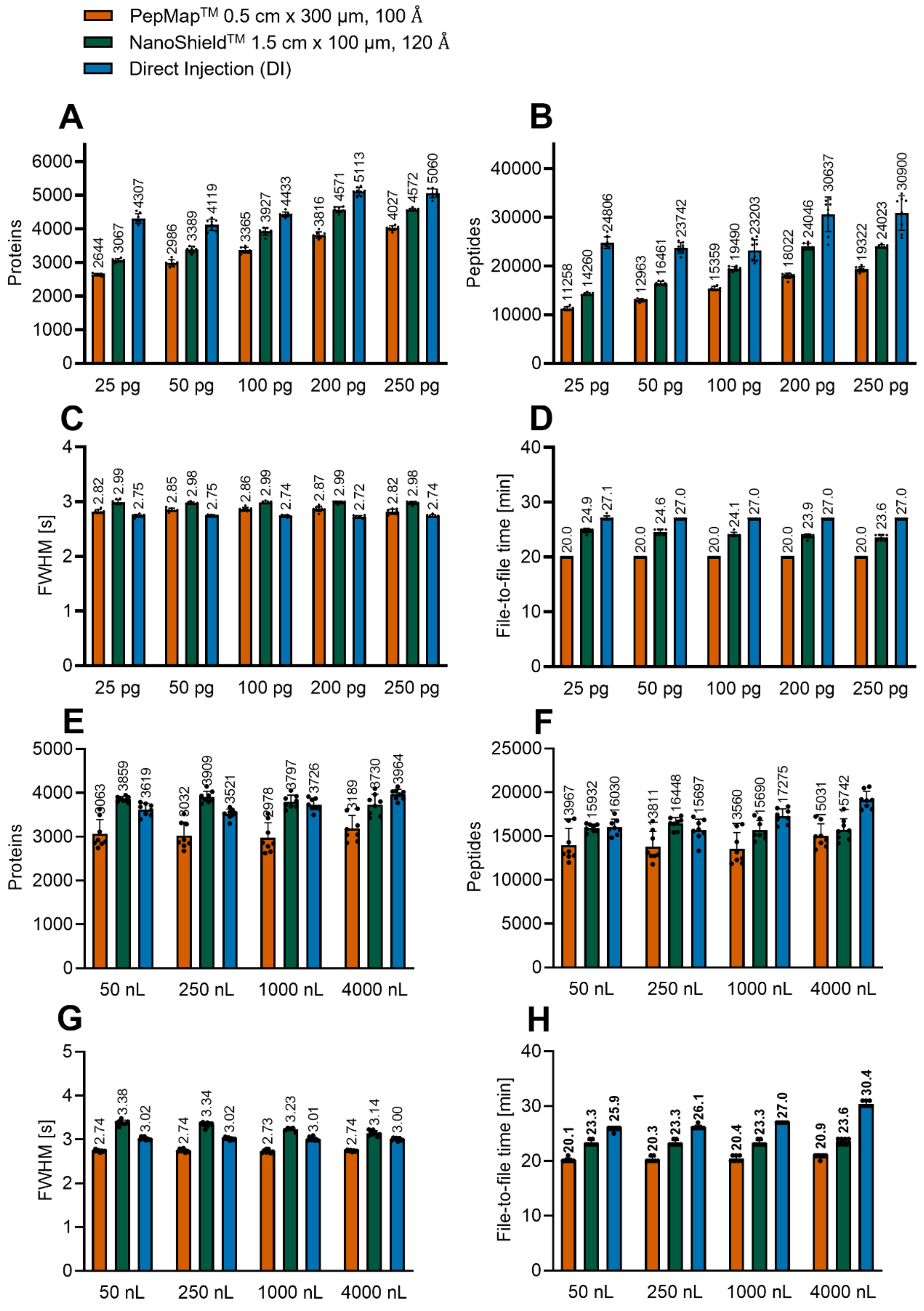

***Supplementary Figure 1. NanoShield™ robustness across sample input amount and injection volume (datasets e, f).***

*(A, B) Number of identified proteins (A, 1% FDR at protein level) and peptides (B, 1% FDR at precursor level) for the PepMap™ Neo trap cartridge (red) and the NanoShield™ 1.5 cm trap column (operated in reverse-flush mode, green), and direct injection (DI, blue) across a titration of sample input amount (25, 50, 100, 200, and 250 pg) at a constant injection volume of 1 µL, using a 14 min LC gradient; ; legend dimensions denote packed bed length x inner diameter, stationary phase pore size (dataset e). (C) Average*

*chromatographic peak width (full width half maximum, (FWHM)). (D) File-to-file time, defined as the total elapsed time between successive raw file writes, encompassing sample loading, equilibration, separation, and column wash. (E, F) Number of identified proteins (E) and peptides (F) for the same three conditions across a titration of sample injection volume (50, 250, 1000, and 4000 nL) at a constant sample input amount of 250 pg, using the same 14 min LC gradient (dataset f). (G) FWHM. (H) File-to-file time. 250 pg Pierce™ HeLa digest standard per injection. Data points represent independent sample injections (n = 8 for PepMap and direct injection in panels A–D, except direct injection at 250 pg with n = 7; n = 7 for NanoShield 1.5 cm across all input amounts in panels A–D; n = 8 for all conditions in panels E–H); bars and numbers show the mean; error bars show ± standard deviation (SD). No statistical testing was performed for this data.*

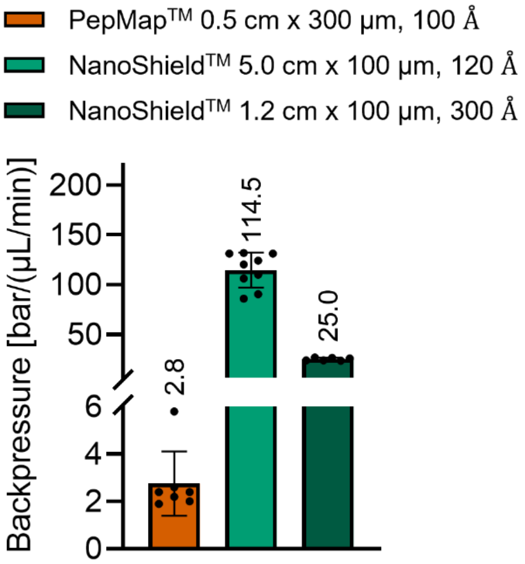

***Supplementary Figure 2. Trap column backpressure.***

*Backpressure for PepMap™ Neo trap cartridge (red), NanoShield™ 5 cm (light green), and NanoShield™ 1.2 cm trap column (dark green), recorded using the trap-column-only resistance test, D01 script on a Vanquish Neo UHPL; legend dimensions denote packed bed length x inner diameter, stationary phase pore size. Data points represent independent measurements across at least 6 instances per type; bars and numbers show the mean; error bars show ± SD. Note the split y-axis to accommodate the large range in recorded backpressure values.*

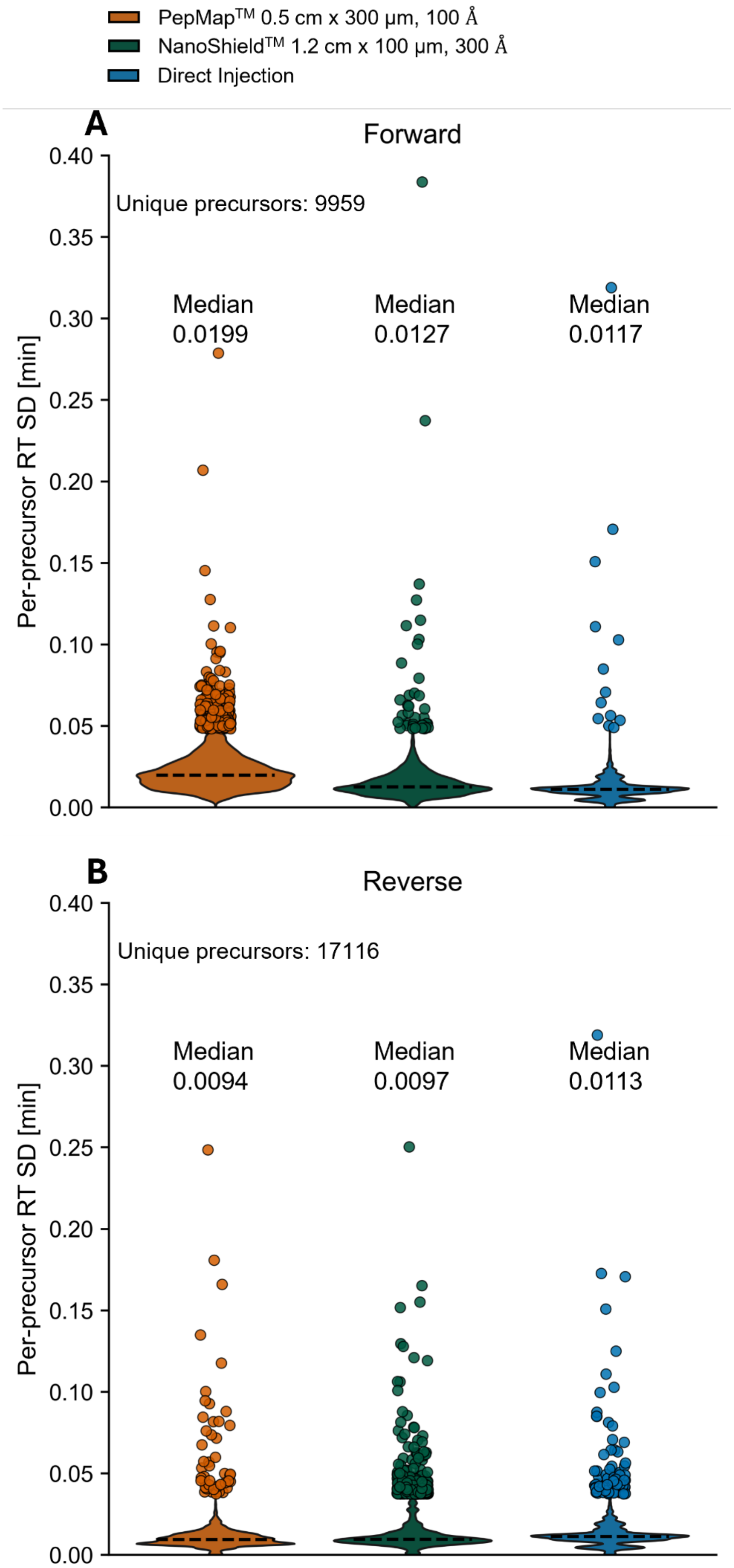
PepMapTM 0.5 cm x 300 µm, 100 Å
NanoShieldTM 1.2 cm x 100 µm, 300 Å
Direct Injection
A
Forward
Unique precursors: 9959
Median 0.0199
Median 0.0127
Median 0.0117
Per-precursor RT SD [min]
0.40
0.35
0.30
0.25
0.20
0.15
0.10
0.05
0.00
B
Reverse
Unique precursors: 17116
Median 0.0094
Median 0.0097
Median 0.0113
Per-precursor RT SD [min]

***Supplementary Figure 3. Retention time (RT) stability across trap columns and direct injection in forward- and reverse-flush mode (dataset g).***
*Per-precursor RT stability for PepMap™ Neo trap cartridge (red), NanoShield™ 1.2 cm trap column (green), and direct injection (blue), operated in forward-flush (A) and reverse-flush (B) mode, using a 14 min LC gradient; legend dimensions denote packed bed length x inner diameter, stationary phase pore size. For each condition, precursors identified in every replicate of every condition were retained, and the SD of RT across the 6 technical replicates was calculated per precursor; violin plots show the distribution of these per-precursor RT SD values, with the median indicated above each violin. Precursors with an RT SD exceeding the plotted range are shown as individual jittered points. Data are derived from the same runs and processing as Figure 4.*

***Supplementary Table 1. Manufacturer-reported specifications of the evaluated trap column formats.*** *Particle and pore size, inner diameter, packed bed length, pressure rating, and loading capacity are listed for the NanoShield™ trap columns (IonOpticks) and the PepMap™ Neo trap cartridge (Thermo Fisher Scientific).*

| Specification | NanoShield™ 1.2 cm x 100 µm, 300 Å | NanoShield™ 1.5/1.2 cm x 100 µm, 300 Å | NanoShield™ 5 cm x 100 µm, 120 Å | PepMap™ Neo Trap Cartridge |
|---|---|---|---|---|
| Stationary phase - particle size | 3 µm | 3 µm | 3 µm | 5 µm |
| Stationary phase - pore size | 300 Å | 120 Å | 120 Å | 100 Å |
| Dimensions - inner diameter | 100 µm | 100 µm | 100 µm | 300 µm |
| Dimensions - packed bed length | 1 cm | 1.2 cm | 5 cm | 0.5 cm |
| Pressure rating | 1500 bar | 1500 bar | 1500 bar (rec.: < 1000 bar) | 1500 bar |
| loading capacity | 200 - 250 ng | 200 – 250 ng | 1 µg | - |

#### DATASET OVERVIEW

Seven datasets (a–g), each corresponding to a distinct experimental comparison presented in the main text or supplementary figures, were acquired over the course of this study. Trap column prototypes evaluated within the same dataset were run back-to-back under matched conditions; experimental details specific to each dataset are summarized below.

**Dataset a**. Acquired on the timsTOF Ultra 2, comparing the NanoShield™ 5 cm x 100 µm trap column against the PepMap™ Neo trap cartridge, both operated in reverse-flush mode, across two LC gradient lengths (14 min and 6.75 min), using an Aurora Rapid CSI 4th Gen 8 cm x 75 µm analytical column. 250 pg Pierce™ HeLa digest standard was injected per run at a sample injection volume of 50 nL.

**Dataset b**. Acquired on the Orbitrap Astral, comparing the NanoShield™ 5 cm x 100 µm trap column against the PepMap™ Neo trap cartridge, both operated in reverse-flush mode, across two LC gradient lengths (25 min and 6.25 min), using an Aurora Rapid XT 4th Gen 8 cm x 75 µm analytical column. 250 pg Pierce™ HeLa digest standard was injected per run at a sample injection volume of 50 nL.

**Dataset c**. Acquired on the Orbitrap Astral, comparing the PepMap™ Neo trap cartridge, the NanoShield™ 5 cm x 100 µm trap column, and the NanoShield™ 1.2 cm x 100 µm trap column, each operated in both forward- and reverse-flush mode, using a 14 min LC gradient and an Aurora Elite XT 4th Gen 15 cm x 75 µm analytical column. 250 pg HeLa digest standard was injected per run at a sample injection volume of 50 nL.

**Dataset d**. Acquired on the Orbitrap Astral, comparing the NanoShield™ 1.2 cm x 100 µm trap column, operated in reverse-flush mode, against direct injection, using the same 14 min LC gradient and an Aurora Elite XT 4th Gen 15 cm x 75 µm analytical column. 250 pg Pierce™ HeLa digest standard was injected per run at a sample injection volume of 50 nL.

**Dataset e**. Acquired on the Orbitrap Astral, comparing the PepMap™ Neo trap cartridge, the NanoShield™ 1.5 cm x 100 µm trap column (reverse-flush mode), and direct injection across a titration of sample (Pierce™ HeLa digest standard) input amount (25–250 pg) at a constant injection volume of 1 µL, using a 14 min LC gradient and an Aurora Elite XT 4th Gen 15 cm x 75 µm analytical column.

**Dataset f**. Acquired on the Orbitrap Astral, comparing the PepMap™ Neo trap cartridge, the NanoShield™ 1.5 cm x 100 µm trap column (reverse-flush mode), and direct injection across a titration of sample injection volume (50–4000 nL) at a constant sample input amount of 250 pg Pierce™ HeLa digest standard, using the same 14 min LC gradient and an Aurora Elite XT 4th Gen 15 cm x 75 µm analytical column.

**Dataset g**. Acquired on the Orbitrap Astral, comparing the PepMap™ Neo trap cartridge and the next-generation NanoShield™ 1.2 cm x 100 µm trap column (300 Å stationary phase), each operated in both forward- and reverse-flush mode, against direct injection, using a 14 min LC gradient and an Aurora Rapid TS 3rd Gen 8 cm x 75 µm analytical column. 250 pg Pierce™ HeLa digest standard was injected per run at a sample injection volume of 50 nL.

### METHODS

#### *Sample preparation and liquid handling*

For all samples lyophilized Pierce™ HeLa protein digest standard (Thermo Fisher Scientifc, catalog nr. 88328) was reconstituted by adding 100 µL of 0.1% TFA and 0.015% DDM in Milli-Q water and 10 s vigorous vortexing. Required dilutions for different sample amounts and sample volumes of this HeLa digest for direct injection or trap-and-elute workflows were either prepared by manual pipetting in polypropylene HPLC vials (Waters, catalog nr. 186009186) capped with preslit PTFE/silicon septum (Waters, catalog nr. 186005827) or using 384 well plates (Eppendorf, Twin tec PCR plates, catalog nr. 0030128508) with pipetting performed by the Mosquito LV liquid handling robot (SPT Labtech). Plates were sealed with adhesive PCR foil seal (VWR, catalog nr. 732-3218).

#### *LC instrumentation and gradients*

For performing low-nanoflow liquid chromatography a Vanquish Neo UHPLC system (Thermo Fisher Scientific, catalog nr. VN-S10-A-01) coupled to one of the following analytical columns that were used in this work: Aurora Rapid CSI 4th Gen 8 cm x 75 µm (Ion Opticks), Aurora Rapid XT 4th Gen 8 cm x 75 µm (Ion Opticks), Aurora Elite XT 4th Gen 15 cm x 75 µm (Ion Opticks), or Aurora Rapid TS 3rd Gen 8 cm x 75 µm (Ion Opticks). All columns feature an integrated emitter. The operated column was heated to 50 °C using the HeatSync™ column heater (Ion Opticks, catalog nr. HSCOLHTR01) or the integrated column oven of the timsTOF Ultra 2 (Bruker Daltonics). 0.1% FA in Milli-Q water served as mobile phase A and weak wash liquid. 80% acetonitrile and 0.1% FA in Milli-Q water served as mobile phase B and strong wash liquid. The autosampler was operated at a temperature of 7 °C.

For direct injection workflow, samples were loaded at 1000 bar using pressure control mode with the loading volume set to "Automatic", corresponding to 5 µL used to transfer the sample plug to the separation column and the fast loading option turned on. Column wash and equilibration were done in pressure control mode at 1000 bar with an equilibration factor of 3 (used volume = void volume of column x equilibration factor) and fast equilibration turned on.

For trap-and-elute workflow, samples were loaded at 800 bar using pressure control mode with the loading volume set to "Automatic", corresponding to 5 µL used to transfer the sample plug to the separation column and the fast wash and equilibration option for the trap column turned on.

All gradients are described in detail below.

#### 25 min gradient

| Time | Duration [min] | Flow [µL/min] | Percentage B [%] | Volume [µL] |
|---|---|---|---|---|
| | Run | | | |
| 0.0 | 0.0 | 0.45 | 1.0 | 0.00 |
| 0.1 | 0.1 | 0.45 | 4.0 | 0.05 |
| 1.9 | 1.8 | 0.45 | 12.0 | 0.81 |
| 2.0 | 0.1 | 0.2 | | 0.03 |
| 12.0 | 10.0 | 0.2 | 22.5 | 2.00 |
| 19.5 | 7.5 | 0.2 | 40.0 | 1.50 |
| 19.5 | Column Wash | | | |
| 22.0 | 2.5 | 0.3 | 99.0 | 0.63 |
| 25.0 | 3.0 | 0.3 | 99.0 | 0.90 |
| 25.0 | Stop Run | | | |
| 25.0 | Column Equilibration | | | |

#### 14 min gradient – on timsTOF Ultra 2

| Time | Duration [min] | Flow [µL/min] | Percentage B [%] | Volume [µL] |
|---|---|---|---|---|
| | Run | | | |
| 0.000 | 0.0 | 0.80 | 8.0 | 0.00 |
| 0.875 | 0.875 | 0.80 | 16.0 | 0.88 |
| 0.975 | 0.100 | 0.25 | 17.0 | 0.06 |
| 3.000 | 2.025 | 0.25 | 23.0 | 0.51 |
| 10.500 | 7.500 | 0.25 | 35.0 | 1.88 |
| 12.500 | 2.000 | 0.25 | 56.0 | 0.50 |
| 12.500 | Column Wash | | | |
| 13.000 | 0.500 | 1.00 | 99.0 | 0.31 |
| 14.000 | 1.000 | 1.00 | 99.0 | 1.00 |
| 14.000 | Stop Run | | | |
| 14.000 | Column Equilibration | | | |

#### 14 min gradient – on Orbitrap Astral

| Time | Duration [min] | Flow [µL/min] | Percentage B [%] | Volume [µL] |
|---|---|---|---|---|
| | Run | | | |
| 0.0 | 0.0 | 0.80 | 6.0 | 0.00 |
| 0.5 | 0.5 | 0.80 | 10.0 | 0.40 |
| 1.0 | 0.5 | 0.25 | 12.0 | 0.26 |
| 10.5 | 9.5 | 0.25 | 37.0 | 2.38 |

| | | | | |
|---|---|---|---|---|
| 12.5 | 2.0 | 0.25 | 56.0 | 0.50 |
| 12.5 | Column Wash | | | |
| 13.0 | 0.5 | 0.8 | 99.0 | 0.26 |
| 14.0 | 1.0 | 0.8 | 99.0 | 0.80 |
| 14.0 | Stop Run | | | |
| 14.0 | Column Equilibration | | | |

##### 6.75 min gradient – on timsTOF Ultra 2

| Time | Duration [min] | Flow [µL/min] | Percentage B [%] | Volume [µL] |
|---|---|---|---|---|
| | Run | | | |
| 0.0 | 0.0 | 0.80 | 5.0 | 0.00 |
| 5.25 | 5.25 | 0.80 | 35.0 | 4.20 |
| 5.25 | Column Wash | | | |
| 5.75 | 0.50 | 0.80 | 99.0 | 0.40 |
| 6.75 | 1.00 | 0.80 | 99.0 | 0.80 |
| 6.75 | Stop Run | | | |
| 6.75 | Column Equilibration | | | |

##### 6.25 min gradient – on Orbitrap Astral

| Time | Duration [min] | Flow [µL/min] | Percentage B [%] | Volume [µL] |
|---|---|---|---|---|
| | Run | | | |
| 0.0 | 0.0 | 0.80 | 5.0 | 0.00 |

| | | | | |
|---|---|---|---|---|
| 5.250 | 5.250 | 0.80 | 35.0 | 4.20 |
| 5.250 | Column Wash | | | |
| 5.751 | 0.501 | 0.80 | 90.0 | 0.40 |
| 6.252 | 0.501 | 0.80 | 90.0 | 0.40 |
| 6.252 | Stop Run | | | |
| 6.252 | Column Equilibration | | | |

#### *MS instrumentation and data acquisition*

##### Data acquisition – timsTOF Ultra 2

Data acquisition on the trapped ion mobility spectrometry (tims) time-of-flight mass spectrometer (TOF) Ultra 2 (Bruker Daltonics) was performed using dia-parallel accumulation–serial fragmentation (diaPASEF) in positive ion mode. The mass spectrometer was equipped with a captive spray Ultra 2 ion source. Capillary voltage was set to 1500 V with a flow rate for dry gas at 3.0 L/min and a dry gas temperature set to 200 °C. MS scan range was set from 100 to 1700 *m/z*. TIMS was operated in custom mode with a 1/K0 range of 0.64–1.45 V·s/cm², a ramp time and accumulation time of 130 ms each, resulting in a ramp rate of 7.35 Hz at 100% duty cycle. diaPASEF settings included a mass range of 325–1300 *m/z* and an ion mobility range of 0.64–1.45 1/K0. The diaPASEF scan scheme consisted of one MS1 survey scan followed by four MS/MS scans (ramps) per acquisition cycle, with a total of 12 isolation windows distributed across the four MS/MS ramps (three windows per ramp). The precursor isolation windows were arranged diagonally across the *m/z*–ion mobility plane to match the correlation between precursor *m/z* and ion mobility, thereby maximizing the efficiency of precursor sampling within the specified acquisition range. The total cycle time for one complete round of MS1 and diaPASEF MS/MS scans was approximately 0.68 s. Collision energy was applied in a linear, ion-mobility-dependent manner, ramped from 20 eV at 1/K0 = 0.60 V·s/cm² to 59 eV at 1/K0 = 1.60 V·s/cm².

#### Data acquisition – Orbitrap Astral

Data acquisition on the Orbitrap Astral MS (Thermo Fisher Scientific) equipped with a FAIMS Pro Duo interface (Thermo Fisher Scientific) and an EASY-Spray source was performed in DIA using positive ion mode. Spray voltage was applied statically at 2000 V, the ion transfer tube temperature was set to 280 °C, and FAIMS was operated in user-defined mode with an inner electrode temperature of 100 °C, an outer electrode temperature of 80 °C, and a compensation voltage of −48 V. MS1 spectra were acquired using the Orbitrap analyzer at a resolution of 240,000 across a scan range of *m/z* 400–800, with a normalized AGC target of 500% (absolute AGC value of 5.000 x e6) and a maximum injection time of 100 ms. For MS2 in DIA mode using the Astral analyzer, the precursor mass range was set to *m/z* 400–800 with automatically determined, non-overlapping isolation windows of *m/z* 20 (window placement optimization enabled; 20 scan events per cycle), a scan range of *m/z* 150–2000, and normalized HCD collision energy of 25%. Precursor ions were accumulated for a maximum injection time of 40 ms with a normalized AGC target of 800% (absolute AGC value of 8.000 x e4). Data were recorded in centroid mode with a loop time of 0.6 s.

##### *Trap column specifications*

Five trap column configurations were evaluated across this study: a NanoShield™ 5 cm x 100 µm trap column (IonOpticks), two short-format NanoShield™ prototypes of nominally 1.5 cm and 1.2 cm x 100 µm packed bed length (IonOpticks), a next-generation short-format NanoShield™ prototype of nominally 1.2 cm x 100 µm packed bed length featuring a new, larger-pore-size stationary phase (IonOpticks; referred to throughout as "300 Å" to reflect its stationary phase pore size), and the PepMap™ Neo trap cartridge, 0.5 cm x 300 µm (Thermo Fisher Scientific, catalog nr. 174500). Full specifications for all trap columns, including stationary phase particle size and pore size, internal diameter, packed bed length, pressure rating, and loading capacity, are provided in Supplementary Table 1.

The two short-format NanoShield™ prototypes (nominally 1.5 cm and 1.2 cm) are functionally equivalent and differ only in packed bed length, which falls within the manufacturer-specified manufacturing tolerance of ±0.2 cm for these prototype trap columns; both feature the same 120 Å stationary phase, particle size, and internal diameter, and are treated as interchangeable throughout this study. The next-generation short-format NanoShield™ trap column features a distinct stationary phase with a substantially larger pore size (300 Å, compared to 120 Å for the original NanoShield™ stationary phase and 100 Å for the PepMap™ Neo trap cartridge) while otherwise retaining the same particle size, internal diameter, and pressure rating as the preceding short-format NanoShield™ generation.

Unlike the 5 cm NanoShield™ trap column and the PepMap™ Neo trap cartridge, both of which are constructed with symmetric frits at either end and are therefore operationally direction-agnostic, the short-format NanoShield™ trap columns, including the 300 Å stationary phase variant, are manufactured with asymmetric frits of differing length at each end, conferring a manufacturer-specified preferred direction of operation for peptide elution. This preferred direction is indicated on the trap column label by directional arrows, although the column remains physically operable in either orientation.

##### *Backpressure measurements and monitoring*

Trap column back pressure was recorded for each trap column configuration using the D01 diagnostic script of the Vanquish Neo UHPLC system (Thermo Fisher Scientific), which is designed to assess the resistance of individual fluidic path segments of the LC system. The D01 script can be run in full or restricted to the trap column flow path only, by enabling the "Test Only Trap Column Resistance" option; back pressure values reported in this study correspond to this trap-column-restricted test mode. Trap column back pressure, reported in bar/(µL/min), was recorded at least once for each newly mounted trap column, for every trap column type and configuration considered in this study, enabling comparison of the resistance of the different trap column types and tracking of individual trap column condition over time.

##### *Data analysis*

All raw files were processed using DIA-NN[1] (v. 2.7.0 Academia) in library-free mode using an in silico predicted spectral library generated from the UniProt reference human proteome (downloaded 2025-07-30) and from a contaminants database derived from the Cambridge Centre for Proteomics (CCP)[1], the latter being included via the built-in DIA-NN "Contaminants" option. Protease specificity was set to Trypsin/P with a maximum of one missed cleavage, no semi-specific cleavage, and no variable modifications beyond N-terminal methionine excision; carbamidomethylation of cysteine was set as a fixed modification. Peptide length was restricted to 7–30 residues, precursor *m/z* to 300–1800, and fragment ion *m/z* to 200–1800. Protein inference was enabled, scoring was performed in Generic mode, proteotypicity was assessed at the gene level, and machine learning was performed using neural networks with cross-validation. Quantification was performed using the QuantUMS (precision) strategy, and spectral libraries were generated/refined from each DIA-NN search run including IDs, RT, and ion mobility (where applicable) profiling. Speed settings were kept at "Optimal results" for peak filtering and "Balanced" for RT/ion mobility filtering. A precursor and protein group FDR threshold of 1% was applied in all searches. For each dataset, search settings were defined via a DIA-NN pipeline settings file (.pipeline.json) to ensure identical parameterization across all runs of that dataset. Minor instrument-specific adjustments to these default settings, including the generation of an empirical calibration library for the timsTOF Ultra 2 data, are described separately below for each mass spectrometry platform.

#### Data analysis – timsTOF Ultra 2

For data acquired on the timsTOF Ultra 2, raw files were searched using the InfinDIA pre-search mode in DIA-NN. Because InfinDIA pre-search requires an empirical, instrument- and acquisition-specific calibration library rather than relying solely on the in silico predicted library, such a calibration library was first generated for each dataset using DIA-NN's built-in "Create Calibration/Tuning Library (InfinDIA)" wizard, with three representative raw files from the corresponding dataset provided as input; the settings populated by this wizard were likewise saved as a dedicated calibration pipeline settings file to ensure that the calibration step was performed consistently and reproducibly for each dataset. The resulting empirical calibration library was then loaded into a separate, dedicated main-search pipeline settings file and used for calibration during the subsequent InfinDIA main search of the full dataset. For the main search, MS1 and MS2 mass accuracies were set to 15 ppm, calibration accuracy and scan window were left at automatic (0) assignment by DIA-NN, and match-between-runs was disabled. Precursor charge states from 2 to 4 were considered.

#### Data analysis – Orbitrap Astral

For data acquired on the Orbitrap Astral, raw files were searched using the "Prediction from FASTA" mode in DIA-NN, with MS1 accuracy, MS2 accuracy, calibration accuracy, and scan window all left at automatic (0) assignment by DIA-NN, cross-run normalization disabled, and match-between-runs disabled. Precursor charge states from 1 to 4 were considered. As for the timsTOF Ultra 2 data, search settings for each dataset were defined via a dedicated DIA-NN pipeline settings file to ensure consistent parameterization across all runs of that dataset.

##### *Post-processing*

All post-processing of DIA-NN output was performed in Python (v3.10.12) using custom scripts built on the pandas, NumPy, and itertools libraries for data handling and calculations, scikit-learn for isotonic regression, and matplotlib and seaborn for visualization. DIA-NN's main report output (report.parquet) was read into pandas dataframes and, where relevant, split by individual run for downstream processing.

For the determination of protein group, protein, and precursor identification numbers, the DIA-NN main report was filtered independently at each respective level: protein groups were filtered at a protein group Q-value of ≤0.01 (PG.Q.Value), proteins at a protein Q-value of ≤0.01 (Protein.Q.Value), and precursors at a Q-value of ≤0.01 (Q.Value), with only proteotypic entries (Proteotypic = 1) retained at each level. Peptide numbers were derived from this proteotypic, FDR-filtered precursor table by counting unique modified sequences (Modified.Sequence), collapsing precursors differing only in charge state into a single peptide entry, whereas unique precursor counts (Precursor.Id) retained differently charged precursors of the same sequence as distinct entities. Average chromatographic peak width was calculated as the mean full width half maximum (FWHM) across all proteotypic, FDR-filtered precursors identified in each run. File-to-file time, defined as the total elapsed time between successive raw file writes and encompassing sample loading, trap column (or analytical column, for direct injection) equilibration, chromatographic separation, and column wash, was extracted from the LC method run logs for each acquisition.

For the number of identified peptides across the LC gradient, the DIA-NN main report was first filtered at a precursor Q-value of ≤0.01 (run-specific), a global precursor Q-value of ≤0.01 (Global.Q.Value), a global protein group Q-value of ≤0.01 (Global.PG.Q.Value), and a quantity quality of >0.5 (Quantity.Quality). The number of identified peptides was determined per condition (trap column type or direct injection) and separately for forward- and reverse-flush mode, using peptides identified in every replicate of that condition. For each peptide retained by this completeness filter, a single representative RT was calculated as the mean RT across all its observations (replicates and charge states) within the condition. Because trap-and-elute workflows introduce additional dead volume relative to direct injection, causing a gradient delay whereby the same peptide elutes later purely due to plumbing rather than differences in peptide hydrophobicity, RTs were normalized across conditions prior to binning. For each trap-column condition, a isotonicc regression was fitted by mapping its peptide RTs onto the direct-injection RT scale, using as anchors the peptides shared between that condition and direct injection; this calibration was then applied to all peptides of the respective condition. Direct injection served as the reference scale. The normalized RT axis was divided into 100 equal-width bins spanning the global normalized RT range across all conditions, and the number of unique identified peptides was counted per condition within each bin, yielding one profile of peptide identifications across the (normalized) gradient per condition. Forward- and reverse-flush modes were processed and binned independently.

For quantitative reproducibility analysis, the DIA-NN main report was first filtered at a precursor Q-value of ≤0.01 (run-specific), a global precursor Q-value of ≤0.01 (Global.Q.Value), a global protein group Q-value of ≤0.01 (Global.PG.Q.Value), and a quantity quality of >0.5 (Quantity.Quality). Runs were assigned to their respective experimental conditions based on acquisition date, and protein group quantities (PG.MaxLFQ) were restricted to protein groups with non-zero, non-missing values. The coefficient of variation (CV) of protein group quantity was then calculated, for each condition, as the ratio of the SD to the mean of PG.MaxLFQ values across replicates.

A completeness filter was applied prior to CV calculation, retaining only protein groups quantified in every replicate of every condition tested, enabling direct comparison of the same protein group set across conditions. Protein groups matching contaminant (cRAP)[2] entries were excluded from all CV analyses prior to export.

For RT stability, the DIA-NN main report was first filtered at a precursor Q-value of ≤0.01 (run-specific), a global precursor Q-value of ≤0.01 (Global.Q.Value), a global protein group Q-value of ≤0.01 (Global.PG.Q.Value), and a quantity quality of >0.5 (Quantity.Quality). RT stability was assessed per precursor (defined as the combination of peptide sequence and charge state, Precursor.Id) within each condition, using only precursors identified in every replicate of every condition tested, ensuring a fully paired design across conditions. For each precursor and condition, the SD of RT across the replicate injections was calculated as a single, independent measure of RT variability.

Troubleshooting and fine-tuning of custom Python scripts used throughout this study was supported by generative artificial intelligence (Gen-AI) using Claude Sonnet 5 in Abacus (abacus.ai). All custom code used for data processing and analysis, implemented as Jupyter notebooks, will be made publicly available on GitHub.

##### *Statistical analysis and data visualization*

Most data visualization was performed using GraphPad Prism (v. 11) based on tables of pre-processed results generated by the custom Python scripts described above. Statistical comparisons between conditions were performed in GraphPad Prism using an unpaired, two-tailed Welch's t test (assuming Gaussian distribution of the underlying data, without assuming equal SD between groups), except for the RT stability analysis described below.

RT stability plots and statistics were generated directly in Python using matplotlib, seaborn, pandas, NumPy, and SciPy. Because the same precursors were present in all three conditions, per-precursor RT variability (SD) was treated as paired, repeated-measures data rather than independent samples, and compared across conditions using the non-parametric Friedman test as the omnibus test, with Kendall's W reported as the effect size; no normality assumption is required for this rank-based test. Where the Friedman test was significant, pairwise comparisons were performed using Wilcoxon signed-rank tests with Holm correction for multiple comparisons, with matched-pairs rank-biserial correlation r reported as the pairwise effect size. RT variability was visualized as violin plots of the per-precursor SD values used in the corresponding statistical tests, with extreme outlier precursors shown as individual jittered scatter points rather than extending the violin tail. All custom Jupyter notebooks used for data processing, analysis, and figure preparation are shared alongside this manuscript (see Post-processing subsection above), providing full detail on all data handling and visualization steps.

### REFERENCES

1. Demichev, V., Messner, C. B., Vernardis, S. I., Lilley, K. S. & Ralser, M. DIA-NN: neural networks and interference correction enable deep proteome coverage in high throughput. *Nat. Methods* **17**, 41–44 (2020).
2. Mellacheruvu, D. *et al.* The CRAPome: a contaminant repository for affinity purification–mass spectrometry data. *Nat. Methods* **10**, 730–736 (2013).
